# An automated microfluidic platform for toxicity testing based on *Caenorhabditis elegans*

**DOI:** 10.1101/2024.08.27.610021

**Authors:** Jiaying Wu, Jing Xi, Zhouhai Zhu, Wei Zhao, Qiyuan Peng, Chunbo Liu, Ziqian Wan, Yiyi Cao, Xiang Chen, Yang Luan

**Author notes:** Correspondence to: Jing Xi, School of Public Health, Shanghai Jiao tong University School of Medicine, 227 South Chongqing Road, 200025 Shanghai, China., Xiang Chen, National Key Laboratory of Science and Technology on Micro/Nano Fabrication, Department of Micro/Nano Electronics, Shanghai Jiao Tong University, 800 Dongchuan Road, Shanghai, China., Yang Luan, School of Public Health, Shanghai Jiao tong University School of Medicine, 227 South Chongqing Road, 200025 Shanghai, China. Jiaying Wu, Jing Xi and Zhouhai Zhu contributed equally to this work.

## Abstract

Humans are frequently exposed to a multitude of chemicals daily, necessitating efficient methods for rapidly assessing toxicity and potential health risks. Microfluidics has shown promise as an intelligent tool for rapid compound testing, owing to its flexibility in integrating with automated devices. The article introduces an automated microfluidic platform, based on *Caenorhabditis elegans (C. elegans)*, designed for chemical toxicity testing. This platform consists of three modules – worm culture, monitoring, and image analysis – which enable automated worm culturing, drug delivery, periodic monitoring, and automated phenotypic analysis. Researchers have designed a bridged microfluidic chip that permits worms to move freely during experiments and established an economical monitoring module for long-term tracking and periodic imaging. Furthermore, they have developed an automated image analysis algorithm to automatically determine worm bending frequency. The platform was subsequently utilized for long-term toxicological assessments of the organophosphate pesticide and environmental pollutants. Results indicated that the platform can effectively evaluate the general and developmental impacts of chemicals. The automated microfluidic worm analysis platform holds significant potential for applications in drug safety assessmentand drug screening research, contributing to human health and industry advancement.

## Introduction

With the development of technology, large numbers of newly registered chemicals discharge into the environment every day (https://www.cas.org/). Humans are consistently exposed to various chemicals, which demands efficient methods to rapidly evaluate the toxicity and potential health risks to humans. Microfluidics has shown promise as an intelligent tool for rapid testing of compounds due to its flexibility to integrate with automation devices. Current researches based on *in vitro* organ chips or microphysiological systems cannot simulate *in vivo* dynamic process in human bodies^[1, 2]^. *Caenorhabditis elegans (C. elegans)*, a classic model organism, is a powerful alternative *in vivo* assay system that has proven to be a useful tool in toxicology studies^[3, 4]^. Recently, platforms combining microfluidic chip with *C. elegans* are promising for toxicity testing of numerous chemicals^[5, 6]^.

Numerous researchers have developed microfluidic platforms for short-term observation of worms in acute toxicity testing assays^[7, 8]^. These platforms exhibit significant benefits, including minimal reagent usage and the capacity for seamless integration with automated systems. The microchambers can be configured to prevent the overlap of worms along the Z-axis, thereby simplifying the process of automated imaging and monitoring throughout the experiments. In chronic toxicity studies, periodic observation and phenotype analysis are needed to provide information on the cumulative effect of toxins, thus demanding long-term cultivation with periodic drug delivery and medium replenishment. Sahand *et al*. developed a platform for age synchronization and analyzing subtle aging phenotypes of worms, on which worms were fed manually outside and transferred daily into the chip for progeny filtration^[9]^. Researchers have reported the development of platforms capable of sustaining the culture of worms and conducting time-lapse imaging over extended periods within microfluidic chips^[10-12]^. More recently, Le *et al*. introduced a microfluidics-based automated system known as HeALTH, designed to facilitate the lifelong cultivation and behavioral monitoring of worms^[13]^. Traditionally, the observation and analysis of phenotypic characteristics have involved immobilizing worms within microfluidic chips^[9, 10, 14]^. However, there has been a growing trend towards the creation of chips with specialized designs that permit the free movement of worms, thereby enabling the observation of behavioral phenotypes in their entirety. To this end, researchers have also devised algorithms capable of automatically identifying, tracking, and analyzing the behavioral traits of worms^[15-18]^. Despite these advancements, the approaches necessitate substantial computing resources and are computationally intensive for standard computers. In recent times, artificial intelligence (AI) has been integrated into the image processing of worms^[19]^, yet there are limited studies on the use of deep learning for the analysis of worm phenotypes, particularly in images with complex backgrounds such as those from microfluidic chips. In summary, while the aforementioned platforms have made strides in the automation of toxicity testing assays using worms, there remains considerable room for further development.

In this study, we presented a microfluidic platform designed for the automated assessment of chemical toxicity using *C. elegans*, which consists of worm cultivation, drug delivery, regular monitoring and automated phenotype analysis. Initially, we designed a bridge-like microfluidic chip for worm culturing, which allows worms to move freely form larval stage to adult. Additionally, we constructed an economical monitoring module capable of longitudinal tracking and periodic image capture of the worms.. The bend frequency of worms can be automatically determined using an image analysis algorithm previously developed in our study^[20]^. Subsequently, we carried out a whole-life-covered worm culture (17 days) and toxicity evaluation of monocrotophos (MCP) and 2,2’,4,4’-Tetrabromodiphenyl ether (BDE-47) using this microfluidic platform, to validate the capacity of the platform for long-term toxicity testing assays on worms.

## Materials and methods

### Microfluidic device fabrication

The microfluidic chip mold was developed by MEMS (Micro-Electro-Mechanical Systems) process, including metal sputtering and electroplating^[21]^. Liquid PDMS mixture (Sylgard 184 Silicone Elastomer Base : Curing Agent = 10:1) was cast on a silicon mold and cured overnight at 70 °C to fabricate a PDMS chip. The inlet and outlet were punched with needles. Finally, the PDMS chip was bonded to a clean glass by air plasma treatment and stored overnight at 135 °C. For more details, see supplementary materials.

### Worm strains and maintenance

The *C. elegans* strains used in this study were wild-type Bristol N2 , which was obtained from the Caenorhabditis Genetics Center (St. Paul, Minnesota, USA). Nematodes were cultured in the nematode growth medium (NGM, 17 g/L Agar B, 3 g/L NaCl, 2.5 g/L peptone, 25 mM KPO_4_ buffer pH 6.0, 1 mM MgSO_4_, 1 mM CaCl_2_, 5 mg/L cholesterol in ethanol) or in M9 buffer (17 mM K_2_HPO_4_, 42 mM Na_2_HPO_4_, 85 mM NaCl, 1 mM MgSO_4_) with the *E. coli* strain OP50 as nutrient at 20 °C. To obtain a synchronized nematode population, a bleach-sodium hydroxide mixture (0.5 M NaOH and 1% (v/v) NaClO) was used with an established method^[22]^.

### Chemicals

1. Fluoro-20-deoxyuridine (5-FUdR) and dimethyl sulfoxide (DMSO) were purchased from Sigma-Aldrich (St. Louis, Missouri, USA). Monocrotophos (MCP) was purchased from LGC Science (ShangHai Ltd) and dissolved in ultrapure water. 2,2’,4,4’-Tetrabromodiphenyl ether (BDE-47) was obtained from Matrix Scientific (Columbia, South Carolina, USA) and dissolved in DMSO.The stock solutions were subsequently diluted with M9 buffer to obtain the working solutions. Ultrapure water was produced using a Milli-Q system (Millipore, Bedford, MA, USA).

### Lifespan assays

The lifespan assay was performed on age-synchronized adult worms (48 h after hatching) in 96-well plates and PDMS microfluidic chips. Worms were cultured in culturing medium (*E. coli* OP50:M9 buffer 1:200, v/v) and 5-FUdR (final concentration of 20 mg/mL) was added to inhibit reproduction. In the PDMS microfluidic chips, culturing medium was refreshed every day to provide the worms with sufficient nutrients. The worms were stimulated with an acupuncture needle (96-well plates) or by increasing the flow rate (microfluidic chips) every 24 h. If a worm did not respond to any stimulation within 1 min, it was considered dead.

### Developmental toxicity assays

For comparison of the developmental toxicity assays in PDMS microfluidic chips and multi-well plates, N2 worms were treated with MCP (0, 250, 500 μM and 1 mM) and BDE-47 (0, 3, 10 and 30 μg/ml) from the L1 stage (no feeding after hatching). After 24 h, videos of worms in each chamber were recorded and analyzed with automated image analysis module. The same assay was repeated with multi-well plates, and the body length of worms were calculated manually.

### Neurotoxicity assays

Synchronized N2 worms in L4 stage (feeding for 48 h after hatching) were treated with chemicals for 24 h in this assay, and 20 mg/mL 5-FUdR was added to inhibit reproduction. The concentrations of MCP were 0, 250, 500 μM and 1 mM, BDE-47 were 0, 3, 10 and 30 μg/ml. After treatment, worms were rinsed with M9 and recorded by monitoring module. The bend frequencies of worms were calculated with the image analysis module.

### Statistical analysis

GraphPad Prism version 8.0 (GraphPad Software, La Jolla, California, USA) and IBM SPSS Statistics version 25 (SPSS, Chicago, IL, USA) were used for statistical analysis. All data are expressed as mean ± standard error (SE). The log-rank test was used to compare the survival curves of worms cultured on PDMS microfluidic chips and micro-well plates. One-way analysis of variance (one-way ANOVA) followed by the *Dunnett t-*test were used to determine the differences between the treatment groups and control group. A value of *P < 0*.*05* was considered statistically significant.

## Results

### Overview of microfluidic platform for worm toxicity testing

Since current microfluidic platforms for worm toxicity testing usually operate at a low level of automation, we developed an integrated microfluidic platform, which consists of three modules (Fig. 1A). The worm cultivation module includes a microfluidic chip for worm culturing and observation, equipped with pressure-based delivery system. Based on this module, diets and test substances can be automatically added at regular intervals, and worms can be cultured in microfluidic chip until death. During chronic toxicity testing assays, the phenotypes of worms in bright filed can be monitored and collected regularly with the monitoring module, which includes a motion stage and a Pi camera. After capturing images or videos of worms, the body length and bend frequency can be calculated automatically with the image analysis module. Fig.1B shows the physical photograph of the entire platform, which is controlled through the control center.

**Fig 1.**
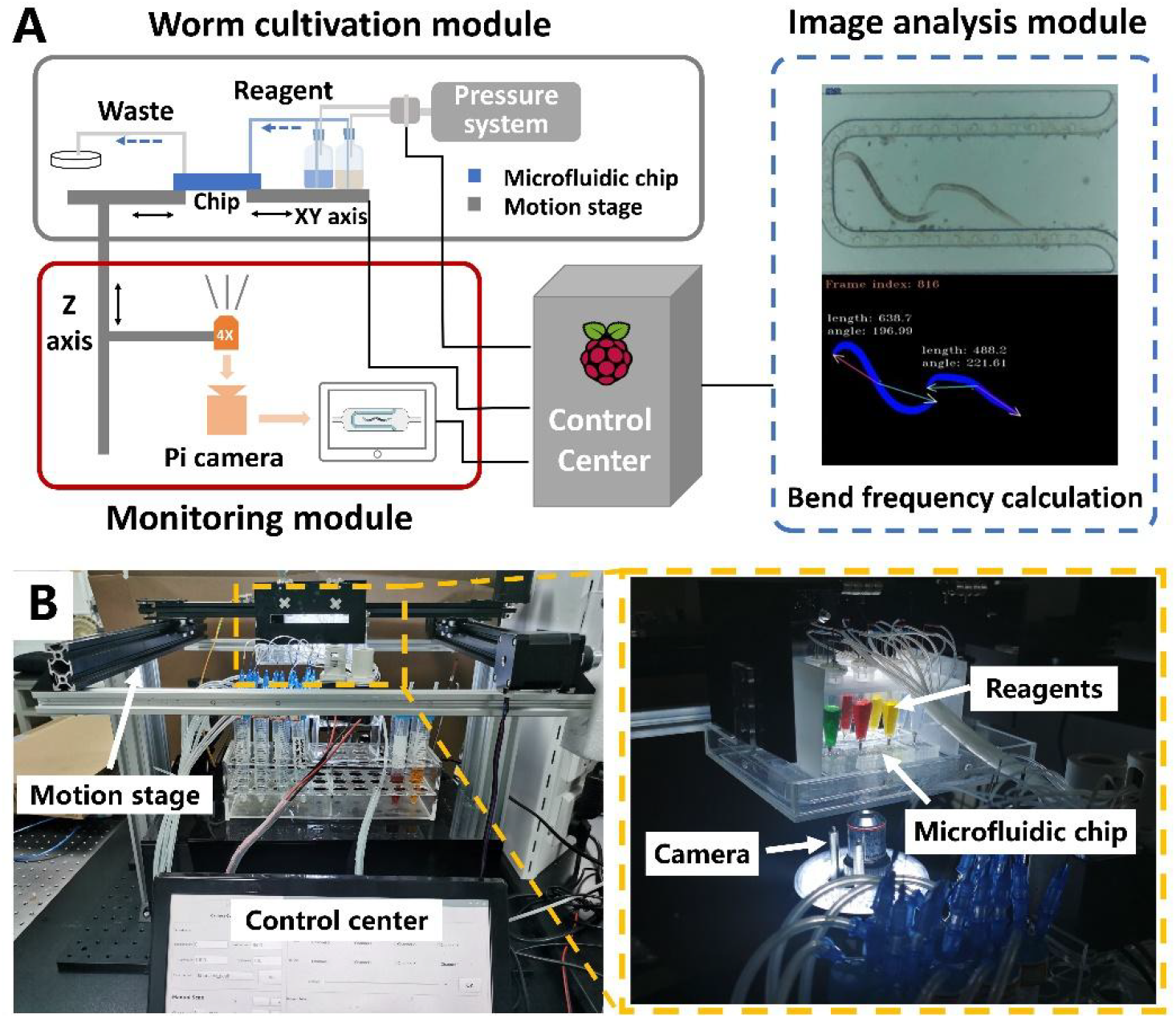
Schematic of the microfluidic platform. **(A)** Diagram of the designed platform consisting of the “worm cultivation module” “monitoring module” and “image analysis module”. **(B)** Physical photograph of the whole platform. Inside the yellow dotted box (graph on the right) is magnified photograph of “worm culturing module”.

### The bridge-like microfluidic chip for worm cultivation

We designed a bridge-like microfluidic chip that allows worms to move freely during the experiment (Fig. 2A). Five parallel chambers were mounted on one chip, and each chamber had a bridge-like structure to allow medium replacement and prevent the worms from escaping. To meet the requirements of worm experiments from the larval stages to adult, the gap should be designed to arrest the worms in L1 stage. Since L1 worms are approximately 10-15 μm wide and sufficiently flexible to squeeze through narrow gaps, an ideal gap should not only arrest the worms but also allow the medium to flow smoothly under relatively low pressure. We tested the working conditions with gap sizes of 2, 5 and 10 μm and confirmed that a 5 μm gap can meet the above requirements (see supplementary materials). To prevent the “bridge deck” from collapsing, pillars were set as support structures at intervals of 100 μm. The designed height of the chambers is 40 μm (approximately equal to the diameter of adult worms) to avoid overlapping more than two worms along the *Z*-axis, which is beneficial for observation and automated analysis. Five inlets and outlets are merged on each chip (Fig. 2C).

**Fig 2.**
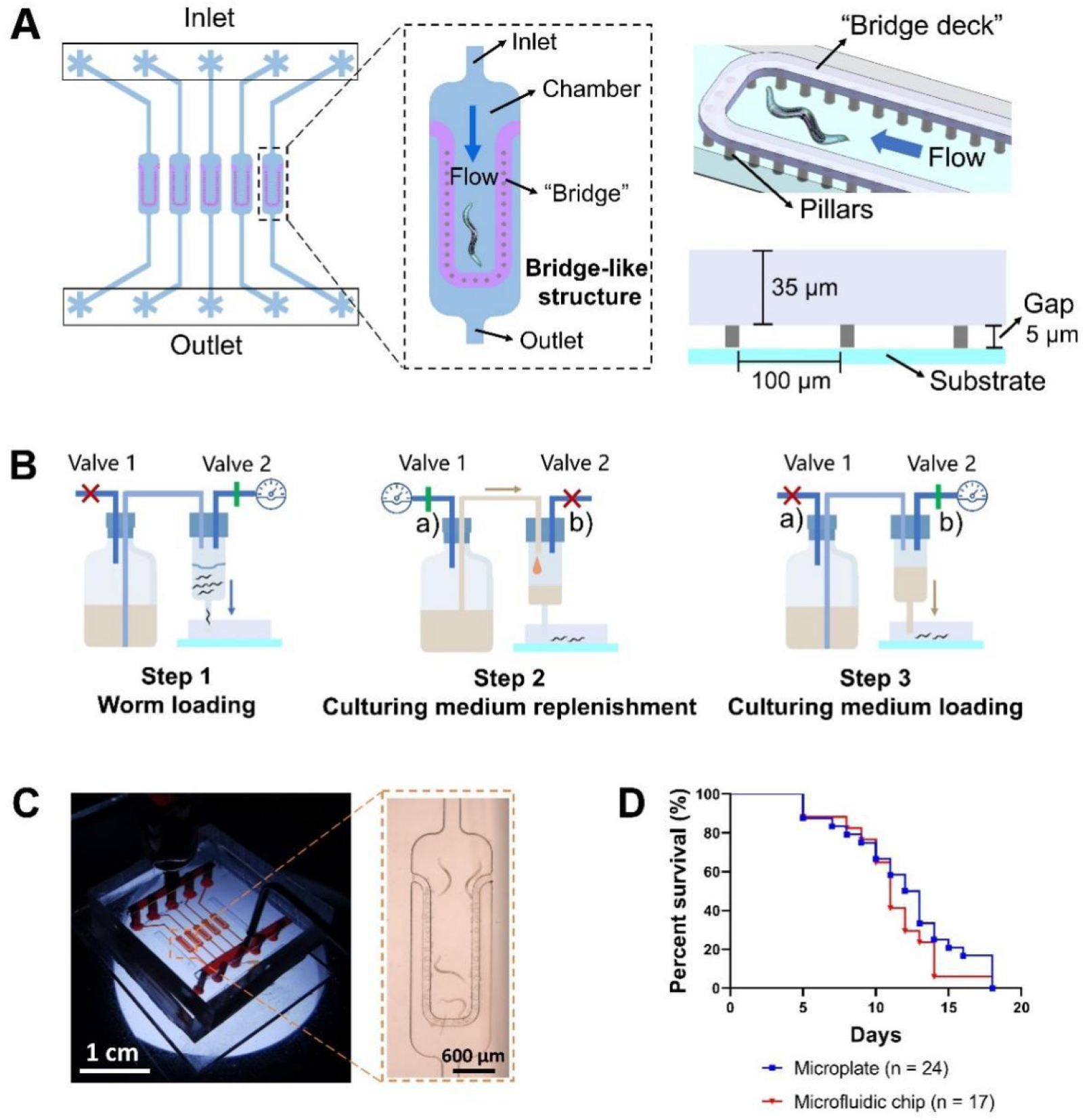
Design and operation of the microfluidic chip. **(A)** Five structures are paralleled in one chip. The inlet and outlet are 150 μm wide and the chamber is 40 μm in height. The “bridge deck” has a thickness of 35 μm and a 5 μm gap underneath to prevent the worms from escaping while allowing the culturing medium to flow. Pillars are set as support structures at intervals of 100 μm. Five inlets and outlets are merged and soaked in liquid to avoid water evaporation. **(B) Procedure of worm loading and nutrient replenishment**. Step 1: valves 1 is closed (red bar) and valve 2 is open (green bar), then worm suspension is loaded into the chip. Step 2: when reagents are exhausted, valves 1 is open and valve 2 is closed. Step 3: nutrient is loaded into the chip. **(C) The physical photograph of the microfluidic chip. (D) Comparison of worm lifespan on multi-well plate and PDMS chip**. The survival curves of the two groups have no significant difference (*p* > 0.05).

The operation procedure of the microfluidic chip is demonstrated as Fig. 2B. Firstly, synchronized worm suspension in the reservoir was loaded into microfluidic chip. During the experiment, culturing medium can be replenished and consistently loaded into the chip at a low flow rate, to provide sufficient nutrients for worms.

To validate the biocompatibility of this microfluidic platform, we cultured worms in multi-well plates and PDMS microfluidic chips until death. Briefly, 24 and 17 worms were included in the multi-well plate group and PDMS microfluidic chips. As depicted in Fig. 2D, the growth status of worms cultured on microfluidic platform are consistent with those in traditional multi-well plates. The average lifespan (± SD) of the worms in multi-well plates and microfluidic chips were 12 (± 4) days and 11 (± 3) days, respectively, which revealed no significant difference between the survival curves of two groups (*P > 0*.*05*) (Fig. 2D). The above results indicated that the platform is suitable for worm growth and automated long-term cultivation, which can relive labor and time compared with traditional assays on multi-well plates.

### Low-cost monitoring module for longitudinal observation and image acquisition

The monitoring module is a low-cost optical imaging system that can ensure stability for the long-term culturing of worms. We built a low-cost three-axis displacement platform using aluminum profiles. Cost of the monitoring module was less than US$ 200. The displacement platform was controlled by an Arduino circuit connected to a Raspberry Pi CPU. The GRBL code library was pre-written in Arduino. The stepper motor and its driver were connected. The Raspberry Pi could send the G code to Arduino to control the motion of the stage. The motion accuracy of this module can easily reach 10 µm. We used a 4× lens and a low-cost Raspberry Pi CCD for imaging.

### Automated image analysis module for phenotype analysis of worms

With the large amount of image data generated in toxicity testing assays, automated data processing can significantly improve the efficiency of the experiment. Therefore, we used the automated image analysis algorithm developed in our previous study^[20]^ to analyze the body length and bend frequency of worms. We used deep learning to extract worm contours from videos that contain chamber structure and impurities in the culture environment. The following process is described as before.

### Toxicity evaluation of MCP and BDE-47 on the platform

Based on this microfluidic platform, we chose MCP and BDE-47 to demonstrate the feasibility of the platform for automated toxicity testing of exogenous chemicals. MCP is an organophosphorus pesticide widely used in agricultural production, the residues of which may be detected frequently in food. As a cholinesterase inhibitor acting on an acetylcholine receptor, MCP is highly neurotoxic^[23]^. As a kind of commercial flame retardant, BDE-47 is the most abundant and prevalent polybrominated diphenyl ethers (PBDEs) congeners in the environment, which has been known to introduce neurodevelopmental toxicity^[24]^.

We treated N2 worms with MCP and BDE-47 for 72 h and evaluated the body length and bend frequency of worms at 24 h. Fig. 3A showed that treatment of MCP and BDE-47 induced significant general toxicity in worms on this platform. The body length was used as the toxic endpoints to evaluate the development toxicity. As shown in Fig. 3B&E, MCP and BDE-47 treatment caused dose-dependent decreases (*p* < 0.001), implying evident developmental toxicity to nematodes. Fig. 3C&F showed that the bend frequency of worms decreased significantly (*p* < 0.001) with the increase in dose after 24 h treatment.

**Fig 3.**
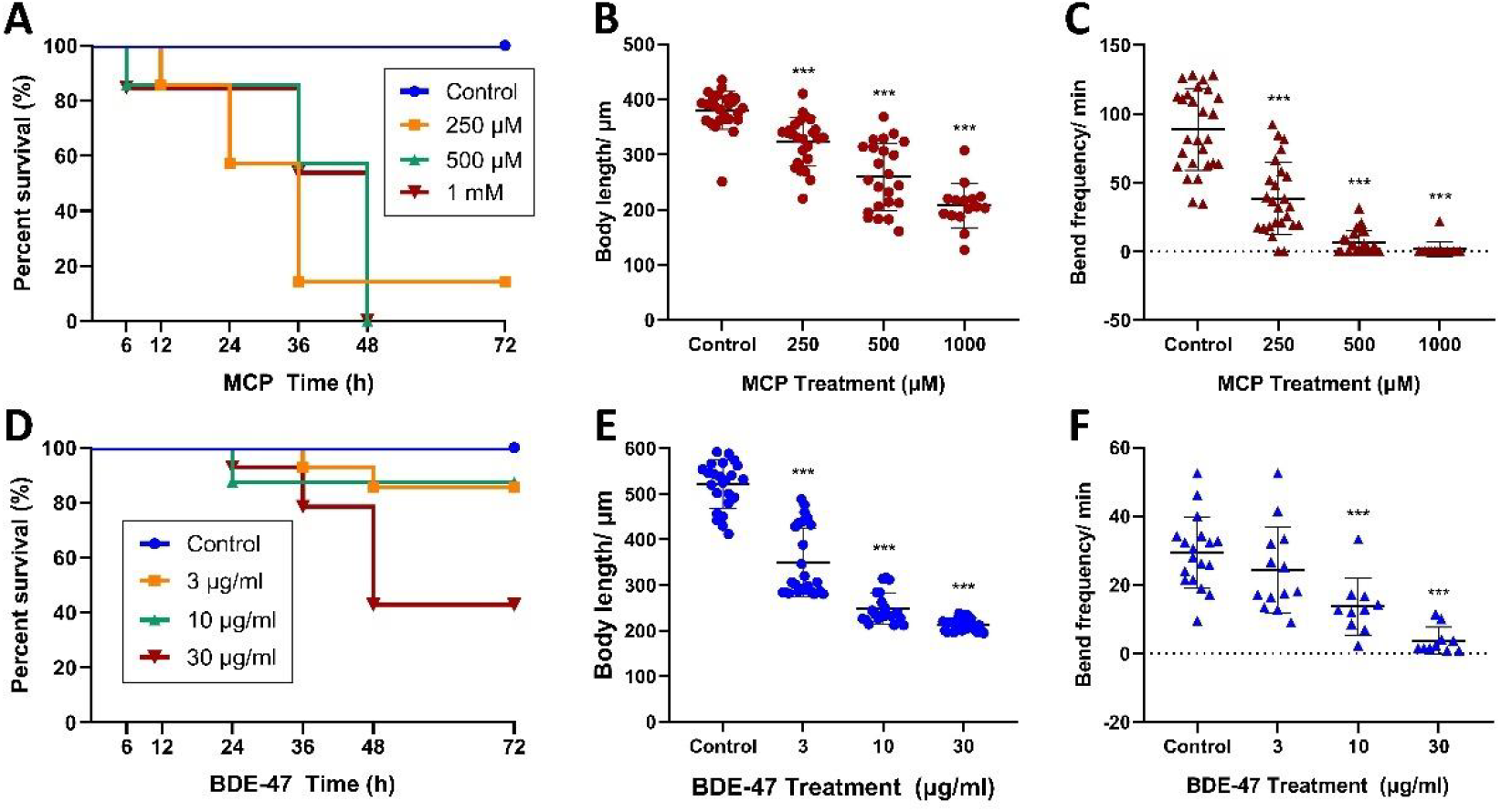
Toxicity evaluation of MCP and BDE-47 on the platform. (A-C) The survival curve, body length and bend frequency of worms treated with MCP. (D-F) The survival curve, body length and bend frequency of worms treated with BDE-47.

The above results demonstrated that this platform can be applied to automated toxicity testing of numerous chemicals that may cause neurotoxicity and development toxicity. In the future, more phenotypes such as brood size and cell apoptosis, can be measured to expand the scope of toxicity evaluation.

## Discussion

With the development of science and technology, microfluidic technology is more and more widely used in the biomedical field. Microfluidic technology has the characteristics of high throughput, low consumption and precise control and is widely used in cell culture, drug screening, disease diagnosis and other fields. As a model organism, *C. elegans* has the characteristics of short life cycle, simple genome and relatively simple nervous system and are ideal models for studying neurotoxicology and neural effects of drugs. In this study, we introduced an automated microfluidic platform based on *C. elegans*, which can realize automation of worm toxicity testing assays including worm culturing, monitoring and phenotype analysis.

Toxicity assays require periodic rinsing and recording of the worms’ phenotypes to evaluate the toxicity of compounds with extended exposure times. In experiments lasting more than 24 hours, the diet-containing culturing medium should also be replenished regularly. Therefore, an automated liquid delivery and switching system is needed. NemaLife has introduced a product that can realize programmable regimens of feeding and reagent addition for toxicity testing (https://www.nemalifeinc.com/infinity-system), but in the latest report, they still use hand-held syringes to manually perform fluid manipulation^[25]^. Some researchers that developed an open-source liquid handling device to realize reagent delivery from large reservoirs to microfluidic chips were faced with the problems of complicated motion platforms and extra on-chip backpressure designs^[26]^. Other researchers applied rotary valves^[27]^ or robotic devices^[28]^ to liquid manipulation, but these fluid switch and delivery systems are limited to complex structures with high-cost, thus limiting the automation of chemical screening. In our study, we designed an automated medium replenishment device to relive the tedious operation of manual liquid change, the cost of which is low enough for laboratories.

Combined with microfluidic technology and nematode model, the neural effects of chemicals or drugs can be studied efficiently, quickly and accurately. Through the platform, the exposure environment of different concentrations of chemicals or drugs to nematodes can be simulated, and the behavioral changes of nematodes, such as movement trajectory, movement speed, reaction time can be observed and recorded. Through the analysis of these data, the neural effects of chemicals or drugs can be evaluated and the scientific basis for the safety assessment can be provided.

In general, the automated microfluidics nematode analysis platform is expected to play a crucial role in the safety assessment of tobacco products, drug screening and neurotoxicology research, contributing greatly to people’s health and the sustainable development of the industry.

## Supplementary Information

Microfluidic device fabrication

Determination of the gap height of the microfluidic chips

## Declarations

### Availability of data and material

The data used and analyzed in the present study are available from the corresponding author on reasonable request.

## Competing interests

The authors declare that they have no competing interests.

## Funding

This work was sponsored by the Open fund of the Joint Institute of Tobacco and Health [grant number 2021539200340048].

## ^1^ Abbreviations

C. elegans: Caenorhabditis elegans;
MCP: monocrotophos;
BDE-47: 2,2’,4,4’-Tetrabromodiphenyl ether;
MEMS: Micro-Electro-Mechanical Systems;
PDMS: Polydimethylsiloxane;
NGM: nematode growth medium;
5-FUdR: 5-Fluoro-20-deoxyuridine;
DMSO: dimethyl sulfoxide;;
SE: standard error;
PBDEs: polybrominated diphenyl ethers;

## Notes

### Competing Interest Statement

The authors have declared no competing interest.

### Summary of Updates

This version of the manuscript updates the author affiliations.

