## Supplemental file for "An automated microfluidic platform for toxicity testing based on *Caenorhabditis elegans*"

*Correspondence to:

**Supplementary Material**

**Microfluidic device fabrication**

**Manufacturing protocol for the microfluidic chip mold:**

(1) Wafer pretreatment: Use a mixed etching solution of hydrogen peroxide and concentrated sulfuric acid to clean a 3-inch monocrystalline silicon wafer for 30 minutes, cool to room temperature, clean the wafer with distilled water in an ultrasonic bath for 5 minutes, and dry it with nitrogen. After the silicon wafer is placed, it is placed in an oven at 180°C for 2 hours to enhance the bonding force between the silicon wafer and the photoresist.

(2) Alloy sputtering: 300 nm thick Cr/Cu electroforming alloy is sputtered on the silicon wafer.

(3) Positive photoresist spin-coated substrate: Pour about 10 ml of AZ 4903 photoresist in the center of the silicon wafer, let it stand for 2 minutes, spin at 200 rpm for 8 seconds and then increase the speed to 2500 rpm for 25 seconds to spread photoresist on the substrate.

(4) Pre-baking: Put the spin-coated silicon wafers in an oven at room temperature, slowly warm up from room temperature to 50 ℃ within 30 minutes, keep the temperature for 1 hour, and then slowly heat up to 90°C within 30 minutes, keep the temperature for 1 hour, then cool to room temperature. The thickness of the positive AZ 4903 photoresist is ~20 μm (measured by a step meter).

(5) Ultraviolet exposure: Use the appropriate mask and expose the photoresist-coated silicon wafer to 365 nm ultraviolet light at 20 mW·cm^-2^ for 60 seconds.

(6) Photoresist development: Use the appropriate developer solution, immerse the wafer in solution, and develop for 1.5 minutes until the pattern is clear.

(7) Electroforming: Put the silicon substrate into the electroplating bath to electroform the copper metallization. The height of the electroplating layer is 5 μm.

(8) Strip photoresist: Wash away the developer on the silicon substrate. Use a series of washes using acetone, ethanol, and distilled water in an ultrasonic bath for 5 minutes each.

(9) Spin-coating negative photoresist: Pour about 10 ml of SU-8 photoresist in the center of the silicon wafer. Let it stand for 2 minutes, spin coat starting at 200 rpm for 8 seconds followed by 2500 rpm for 25 seconds

(10) Pre-baking: Put the spin-coated substrate in an oven, slowly heat it from room temperature to 50 ℃ in 30 minutes and keep it at a constant temperature for 1 hour. Then slowly raise the oven temperature to 90 ℃ within 30 minutes and then hold the temperature for 1 hour. The thickness of the negative SU-8 photoresist is 35 μm (measured by a step meter).

(11) Ultraviolet exposure: Use the appropriate mask and expose for 90 seconds with 365 nm ultraviolet light of 20 mW·cm^-2^.

(12) Development: Use the appropriate developer solution for the negative photoresist. Immerse the wafer in solution and develop for 4 minutes until the pattern is clear.

(13) Mold pretreatment: Place the silicon wafer mold in an oven with a temperature of 80 ℃, keep it at a constant temperature for 1 hour to remove any moisture on the mold, and cool to room temperature. Use 1H, 1H, 2H, 2H-perfluorotrichlorosilane to treat the surface of the silicon wafer mold for 5 minutes to reduce the adhesion between the silicon wafer mold and the PDMS microfluidic chip so that the PDMS polymer can be removed from the mold during peel off.

**Preparation of microfluidic chip**

(1) Fixing: Use double-sided tape to fix the pre-processed chip mold to a 100 mm disposable petri dish to avoid PDMS material from entering the bottom of the silicon wafer mold when PDMS is used for casting.

(2) Prepare PDMS: Use a 10:1 mass ratio of PDMS prepolymer and crosslinking agent. Weigh 20 g of Sylgard 184 prepolymer and 2 g of matching curing agent. Mix the two components thoroughly by stirring. In order to remove bubbles generated during stirring and mixing, which can affect the optical transparency of the device, the mixed PDMS is placed in a vacuum chamber until no bubbles are observed in the PDMS.

(3) Pour PDMS: Use cotton buds to remove dust on the silicon wafer mold. Slowly pour the PDMS material into the silicon wafer mold. Be careful not to introduce bubbles during the process. The thickness of the formed PDMS chip is about 5 mm.

(4) Cure: Place the petri dish with poured PDMS in a 70 ℃ oven for 2-3 hours so that it cures quickly and the structure on the silicon wafer mold is transferred to the PDMS polymer.

(5) Cut and punch: After the PDMS polymer is completely solidified, cool to room temperature. Use a scalpel to cut the chip structure from the mold and carefully peel it off from the silicon wafer mold. Too much force may stress the silicon wafer and cause it to break. Use a spatula to cut the chip according to the designed size and use a No. 19 punch with an inner diameter of 0.67 mm to punch holes at the entrance and exit of the microfluidic chip.

(6) Bonding: Use acetone, ethanol, and distilled water in sequence to clean the glass sheet ultrasonically for 5 minutes. Put the dried glass sheet and the solidified PDMS polymer into a plasma etching machine and treat the PDMS surface and the glass sheet with oxygen plasma for 50 seconds. This process forms a permanent bond by replacing Si-CH_3_ groups with the Si-OH groups.

(7) Place the bonded chips on a hot plate at 135°C for 6 hours to reinforce the bonding effect.

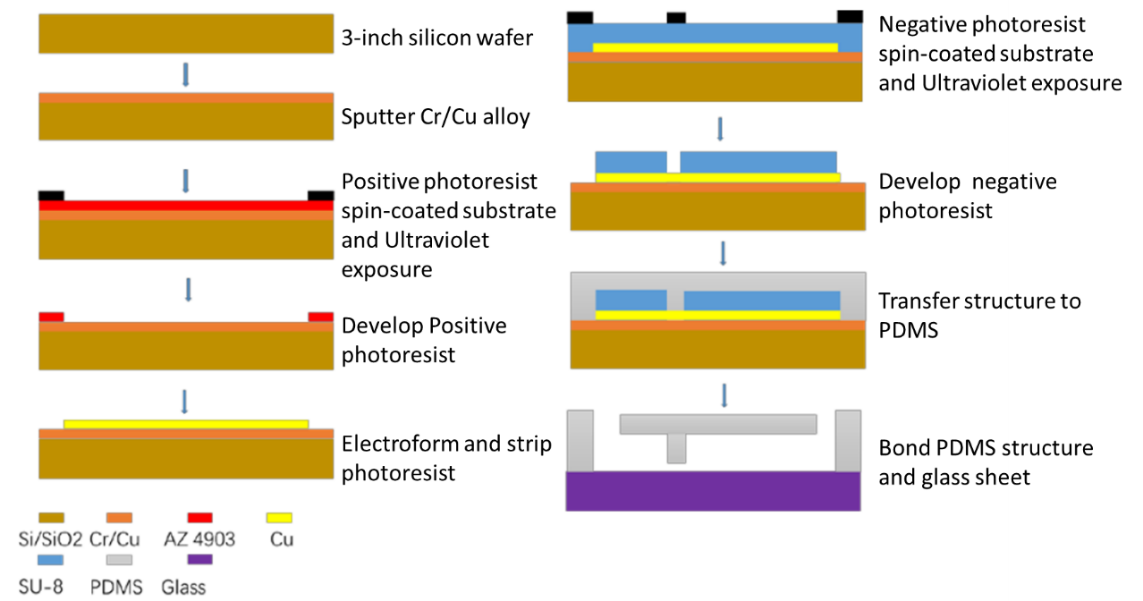

**Fig. S1.** Fabrication process of Microfluidic chip based on MEMS.

**Determination of the gap height of the microfluidic chips**

The height of the gap is a key parameter in microfluidic chip design. To study the effect of the gap height on chip function, we evaluated three chips with gaps of 2, 5, and 10 μm and performed a comparative study.

We used a pressure injection system at the entrance of the microfluidic chip to inject liquid at a pressure of 30 kPa and observed the flow of liquid in the bridge-like structure at gap heights of 2, 5, and 10 μm. With the 10 μm gap, the L1 stage worms easily escaped from the culture chamber because the diameter of the worms in the L1 stage was 10-15 μm; the worms could deform under pressure and escape from the 10 μm gap.

As shown in Fig. S2, in the 2-μm gap structure, the liquid reaches the culture chamber from the entrance of the chip but is enclosed in the bridge-like structure chamber, which makes it difficult for the liquid to flow to the outlet. The gap of 5-μm height can meet the designed chip function and the L1 worms are confined in the chamber for cultivation, but also the metabolic waste of worms and the culture solution containing food can be used by the liquid as media to flow through the gap to the outlet during the worm cultivation process.

The difficulty of the fluid to flow to the outlet in the 2-μm gap experiment is due to the liquid interfacial force. In the process of filling the chip chamber with the culture solution, the two-phase interfacial force between the liquid and the air is an important factor affecting the liquid flow. According to the Young–Laplace equation, the surface pressure difference between the two phases is proportional to the sum of the reciprocal of the chip height *h* and width *w* (Eq. 1).

$\Delta P=2\sigma\left( \frac{1}{h}+\frac{1}{w} \right)$ $\Delta P=2\sigma\left( \frac{1}{h}+\frac{1}{w} \right)$	$\Delta P=2\sigma\left( \frac{1}{h}+\frac{1}{w} \right)$ $\Delta P=2\sigma\left( \frac{1}{h}+\frac{1}{w} \right)$	(1)

In the bridge-like structure, the width *w* of the bridge-like structure is much larger than its height *h*, so the effect of width on the surface pressure difference between the solid and liquid phases is negligible. We only considered the difference in height on the effect of surface pressure difference. The surface tension coefficient of water at 20 °C was 0.0728 N/m. According to Eq. (1), the surface pressure differences between the air and the liquid in the 5 μm and 2 μm structures were 29 kPa and 73 kPa, respectively. Under the same pressure, the bridge-like structure with a height of 2 μm does not facilitate the outflow of the culture fluid in the worm experiment. Although increasing the injection pressure at the inlet can cause the liquid to break through the two-phase interface force and flow to the outlet, it can also delaminate the PDMS chip. For a comprehensive analysis, we chose a 5-μm height difference in the flow channel chamber so that the worms’ metabolic waste in the culture fluid can flow out from the bridge hole of the bridge-like structure, and the worms can be trapped due to the body’s diameter.

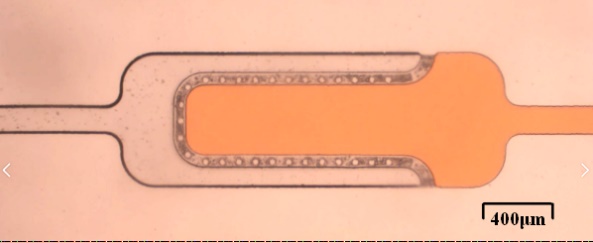

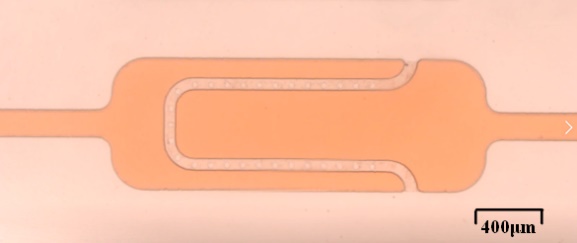

**Fig. S2.** Liquid distribution in the two chambers of 2 μm and 5 μm gaps under the same pressure.
